# Transcriptomically-measured gene expression predicts physiological variation across single neurons in humans and mice

**DOI:** 10.1101/2024.08.26.609746

**Authors:** Rennie M. Kendrick, Scott W. Linderman, Scott F. Owen

## Abstract

Single-cell transcriptomics measures the molecular landscape of individual neurons with unprecedented efficiency and scale. This insight has the potential to advance our understanding of the molecular basis of neuronal function, and to identify druggable targets for disease treatments. However, transcriptomics also suffers from greater measurement noise than traditional techniques (e.g., RT-PCR), which raises questions about its ability to offer insight into function at true single-cell resolution. We tested if transcriptomic data could yield insight into function of individual neurons in human and mouse neocortex by analyzing two datasets collected via Patch-Seq, a powerful technique for obtaining transcriptomic and physiology data from the same neuron. We found that computational models trained on single-cell transcriptomic data robustly predicted physiology of individual neurons. Critically, models trained on single cells outperformed those trained on cell type averages when predicting single-cell physiology.

Thus, the standard approach of denoising single-cell transcriptomic data by averaging on cell types sacrifices functionally-relevant information. Our analysis also revealed novel relationships between gene expression and physiology, including a potential molecular substrate of human-mouse cross-species differences in the speed of single-neuron computation. Broadly, our findings highlight the promise of Patch-Seq for generating new insight into the molecular basis of neuronal function.

## Introduction

Single-cell transcriptomics has revolutionized interrogation of the molecular underpinnings of biological systems. By providing expression of all genes in the genome, this technique has yielded novel insight into the molecular landscapes of individual cells across development^1^ and in response to perturbations, such as in disease.^2,3^ Moreover, the experimentally efficient procedure maximizes data collection from high-value, rare tissues samples, such as from patients.^4^ However, whether transcriptomics can offer meaningful insight into cellular function at true single-cell resolution remains unclear.

First, transcriptomics describes gene expression at the RNA level and, therefore, does not capture many aspects of protein function in the cell. Second, measurement noise—due to dropouts and variable sequencing depths—can limit the signal-to-noise of single-cell transcriptomics relative to previous approaches like bulk RNA sequencing^5^ and RT-PCR.^6,7^

Patch-Seq is one of the only techniques to directly pair transcriptomics with functional measurements from the same cell, and thus offers a unique opportunity to directly test if transcriptomics can predict cellular function at single-cell resolution. Briefly, cellular RNA contents are aspirated and sequenced after whole-cell patch-clamp recordings, enabling a one-to-one mapping of gene expression and physiology.^8–10^

Previous studies have established that ion channel mRNA levels can predict conductance levels of individual channels^11–13^ and variation in intrinsic physiology.^6,7^ However, these studies utilized single-cell RT-PCR after physiology recordings to detect a pre-selected set of ion channel genes, in contrast to the genome-scale expression data afforded by transcriptomics.

We therefore sought to determine if the signal-to-noise of single-cell transcriptomics is sufficient for prediction of intrinsic physiology and discovery of any novel gene expression-physiology relationships.

Recent research has demonstrated that single-cell transcriptomes can predict the synaptic responses^14^ and cellular activation states^15^ of individual neurons. Moreover, Cadwell and colleagues were able to leverage Patch-Seq to predict some physiology features at single-cell resolution in one of the first Patch-Seq datasets ever collected.^8^ However, this sample size was relatively small in the context of linear modeling (58 neurons versus thousands of genes) and therefore necessitated a highly-regularized, sparse model. Moreover, the model did not connect physiology to ion channel gene expression. We set out to extend the scope and interpretability of this work by connecting ion channel gene expression to physiology in recently released larger data sets.^4,16^

A handful of studies have linked transcriptomics to intrinsic physiology using large Patch-Seq datasets. However, these studies averaged gene expression across thousands of cells within each cell type to mitigate measurement noise in gene detection.^9,16–19^ This ‘cell typing’ approach has several limitations: it negates extensive molecular and functional heterogeneity within cell types;^20–28^ it may falsely emphasize cell type marker genes over those that directly drive physiological variation; and it can lead to a ‘failure of averaging’, in which mean values of an input variable (gene expression) map to a different functional output (physiology) than individual samples.^29^

To investigate whether transcriptomic data can predict physiology at single-cell resolution, or if measurement noise obscures biological relationships, we re-analyzed two recently released Patch-Seq datasets from the Allen Institute for Brain Science comprising thousands of mouse interneurons^9^ and hundreds of human pyramidal neurons.^16,30^ We related single-cell transcriptomic data to physiology via well-established, interpretable linear modeling techniques.^31^

Our results reveal that transcriptomically-measured single-cell gene expression can robustly predict single-cell physiology in both of the Patch-Seq datasets that we analyzed. Importantly, the predictive power of these models extended beyond variation across cell types, with models trained on single cells performing better than those trained on the same dataset after being discretized into cell types. This suggests that the signal gained by considering single cell data over cell type averages outweighs any additional noise.

This approach revealed novel gene-physiology relationships in addition to confirming known relationships between ion channel gene expression and physiology. In particular, we uncover a cross-species difference that may be a molecular substrate for the faster kinetics of human relative to mouse neurons that can be tested in follow-up interventional studies.

Furthermore, we find that expression of GPCR and synaptic genes are sufficiently coupled to ion channel gene expression to enable prediction of physiology from these genes as well, suggesting that intrinsic and synaptic physiology are coordinated at the gene expression level. Broadly, our results highlight the power of unbiased, single-cell transcriptomics and its promise for delivering new insight into the molecular basis of cellular function.

## Results

### Gene expression predicts physiology at single-cell resolution

We leveraged an existing mouse interneuron Patch-Seq dataset collected by the Allen Institute^16^ to determine if linear models can predict a set of active and passive physiology features from transcriptomically-measured gene expression at single-cell resolution (Figure 1A). First, we restricted our model to a biologically-relevant gene set (ion channels) to reduce the number of features (genes) relative to samples (cells) and mitigate overfitting. We fit linear models with no regularization (i.e., least squares), L1-regularization (i.e., the LASSO^31^), and L2- regularization (i.e., Ridge^32^). Each model produced largely equivalent performance on held-out data (Supplemental Figure S1). Thus, we proceeded with L1-regularization to take advantage of its built-in tendency to ‘select’ the most useful genes for predicting physiology by dropping unnecessary coefficients exactly to zero.

**Figure 1.**
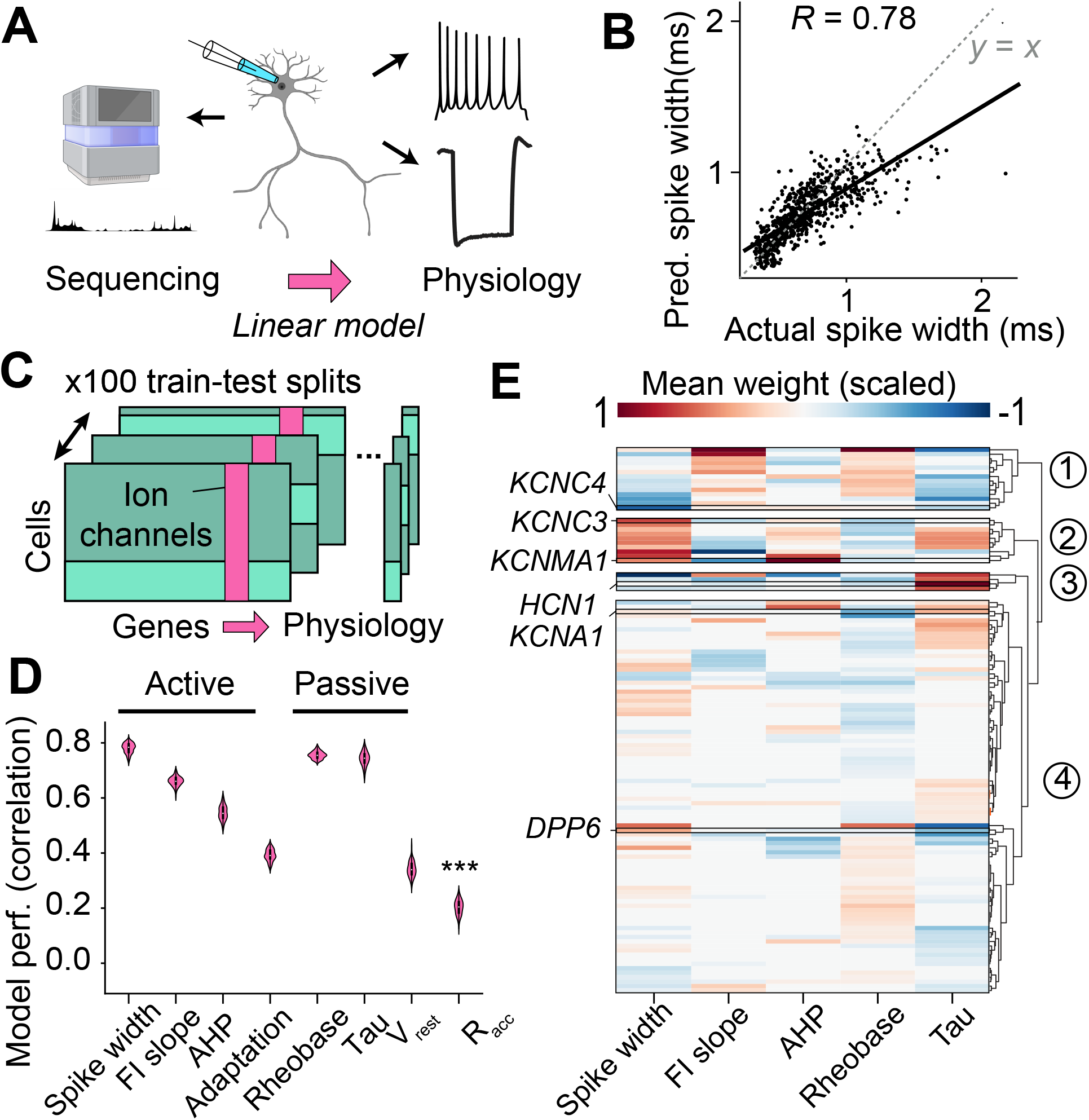
Gene expression predicts physiology at single-cell resolution. A. Published Patch-Seq dataset comprises paired transcriptomics and physiology from single interneurons.^16^ A sparse, L1-regularized linear model was trained to predict intrinsic physiology from ion channel gene expression in single cells. (Neuron created with BioRender.com) **B.** Ion channel gene expression predicts single-cell variation in action potential spike width (Pearson’s *R* = 0.78; *n* = 1396 cells). Dashed line denotes *y*=*x* and displayed data comprises held-out test samples. **C.** Models were trained and tested on 100 random train-test splits of the data to get a distribution of *R*-values. **D.** Active and passive physiology properties can be predicted consistently from ion channel gene expression (*n* = 100 train-test splits) well above and beyond a measure of recording quality, Racc. *** denotes all pairwise *p* < 0.001 on a two-sided unpaired *t*-test with Bonferroni correction. The total number of cells with data for each physiology feature – before dividing into 80/20 train-test splits – is given in Supplemental Table S1. **E.** Model coefficients assigned to ion channel genes for predictive physiology models (mean Pearson’s *R* > 0.5, *n* = 100 train-test splits). Coefficients were scaled to the absolute maximum value for a given physiology model and ion channel genes were clustered according to their assigned coefficients across physiology models.

Remarkably, we found that ion channel gene expression strongly predicted spike width on held-out test samples in a model trained on single-cell data (Figure 1B). This was in direct contrast to the only previous study (to our knowledge) modeling relationships between single-cell physiology and single-cell gene expression collected via Patch-Seq, which could not build a predictive model of spike width.^8^

To test if our results were invariant to the particular train-test split of the data, we refit models to 100 different randomized train-test splits of the data (Figure 1C). We chose this number by increasing the number of train-test splits until further increases failed to reveal additional variability in model performance (Supplemental Figure S1). We found that model performance was consistently high across train-test splits (Figure 1D).

Next, we extended this approach to predict a set of active and passive physiology features. We found that ion channel gene expression predicted physiology accurately for a set of active and passive physiology features (Figure 1D). In addition, ion channel gene expression did not accurately predict access resistance, which is a key technical measure of recording quality that is not biologically driven (Figure 1D). Thus, model predictive accuracy does not depend on artifacts related to data quality.

### Insight into ion channel gene function and model validation

We next mined the resulting model coefficients, considering only models that accurately predicted their respective physiology features (mean Pearson’s *R* > 0.5 measured versus predicted physiology). We had two primary objectives. First, we sought to determine whether gene-physiology relationships were biologically plausible. Second, we aimed to identify potentially novel gene-physiology relationships that may have been missed by previous studies, which either considered fewer ion channels,^6^ genes other than ion channels,^8^ or relied on cell-type averaged gene expression data.^9,16–19^

As expected, the expression of individual ion channel genes often related to multiple different physiology features (pleiotropy).^33,34^ To identify modules of ion channels with shared relationships to one or more physiology features, we clustered channel genes based on their assigned model coefficients across predictive physiology models (Figure 1E; Supplemental Table S2).

Measures of suprathreshold excitability include action potential duration (i.e., spike width) and the relationship between the amount of current injected and the number of elicited spikes (FI slope). Cluster 1 comprised genes whose expression was primarily associated with limiting suprathreshold excitability, including broader action potentials and shallower FI slopes (Figure 1E). The gene encoding Kv3.4 (*KCNC4*) was associated with wider action potentials and a shallower FI slope (Supplemental Table S2; Supplemental Figure S2); this is notable, as prior work describes both positive^35^ and negative^36^ effects of Kv3.4 assembly with other Kv3 channels on spike width.

By contrast, expression of genes in Cluster 2 (Figure 1E) was associated with narrower (i.e., faster) action potentials and a steeper FI slope, which are markers of high-frequency firing. Included in this cluster were genes encoding canonical ion channels critical to fast-spiking, Kv3 subunits (*KCNC1-3*) and Nav1.1 (*SCN1A*). ^6,37–43^ One gene in this cluster, *KCNMA1* (BK channel subunit), was also strongly weighted by a model predicting the membrane potential of the action potential after-hyperpolarization (AHP), but Kv3 subunits were not (Supplemental Figure S2; Supplemental Table S2). This demonstrates that our model can parse subtle mechanistic differences established by previous research in the gating of these channels and their distinctive contributions to action potential phases.^6,44,45^

As for subthreshold excitability, Cluster 3 (Figure 1E) comprised genes that were negatively correlated to the membrane time constant (tau), or that accelerate the return to resting membrane potential after a hyperpolarizing voltage step. This cluster included genes encoding HCN1 (*HCN1*) and Kv7.5 (*KCNQ5*), which underlie h- and M-currents, respectively (Supplemental Table S2; Supplemental Figure S2). Each of these currents can act as ‘voltage clamps’ that oppose voltage deflections^46^, but the Kv7.5 mediated M-current is relatively understudied in cortical interneurons.

Finally, rheobase (the amount of current needed to elicit at least one action potential) was predicted nearly as accurately as spike width and tau, but genes in this model were weighted more diffusely and did not cluster as robustly (Figure 1E). Genes assigned high magnitude coefficients included those encoding Kv1.1 and Kv1.2 voltage-gated potassium channel subunits (*KCNA1* and *KCNA2*), the expression of which was strongly positively correlated to rheobase.^41^ Conversely, expression of the auxiliary subunit *DPP6* was negatively correlated to rheobase.

The expression of *DPP6* and *KCNA1*/*KCNA2* were both negatively correlated to spike width, despite their opposite directions of correlation to rheobase (Supplemental Table S2; Supplemental Figure S2). Mechanistic studies of each of these proteins recapitulate the direction of each of these correlations (*DPP6*^47^; *KCNA1-2*^42,48^). Thus, our model can parse the unique and independent contributions of each ion channel or auxiliary subunit to multiple different physiology features.

Our choice of L1-regularization can enhance model interpretability by ‘selecting’ the most useful genes for predicting physiology and dropping others entirely. However, this can introduce a bias wherein only one gene is retained from a set of correlated genes. As a control, we compared model coefficients from models fit with L1-regularization, with L2-regularization (i.e., Ridge), and no regularization (i.e., least-squares). In general, model coefficients did not vary drastically as a function of regularization method. However, as expected, coefficients fit via L1-regularized linear models were a subset of L2-regularized or unregularized models (Supplemental Figure S1). Moreover, when considering genes that were dropped by our L1- regularized model, but assigned high coefficients by these other approaches, we did not identify additional specific or interpretable gene-physiology relationships (Supplemental Table S3).

### Single-cell gene expression improves model performance relative to cell-type averaging and reveals principles of within-cell-type variation in physiology

Every previous study that has attempted to uncover mechanistically meaningful insight from transcriptomics data has discretized Patch-Seq datasets into cell-type averages. The goal of this approach is to smooth measurement noise by relying on strong covariation between cell type (and subtype) identity, ion channel gene expression (Figure 2A), and physiology^9,16–19^ (Figure 2B). However, this approach squanders the single-cell resolution of Patch-Seq datasets and does not account for extensive within-cell-type variation in physiology.^20–28^ We therefore tested whether models fit to single-cell gene expression data were overall more predictive than those fit to cell type averages.

**Figure 2.**
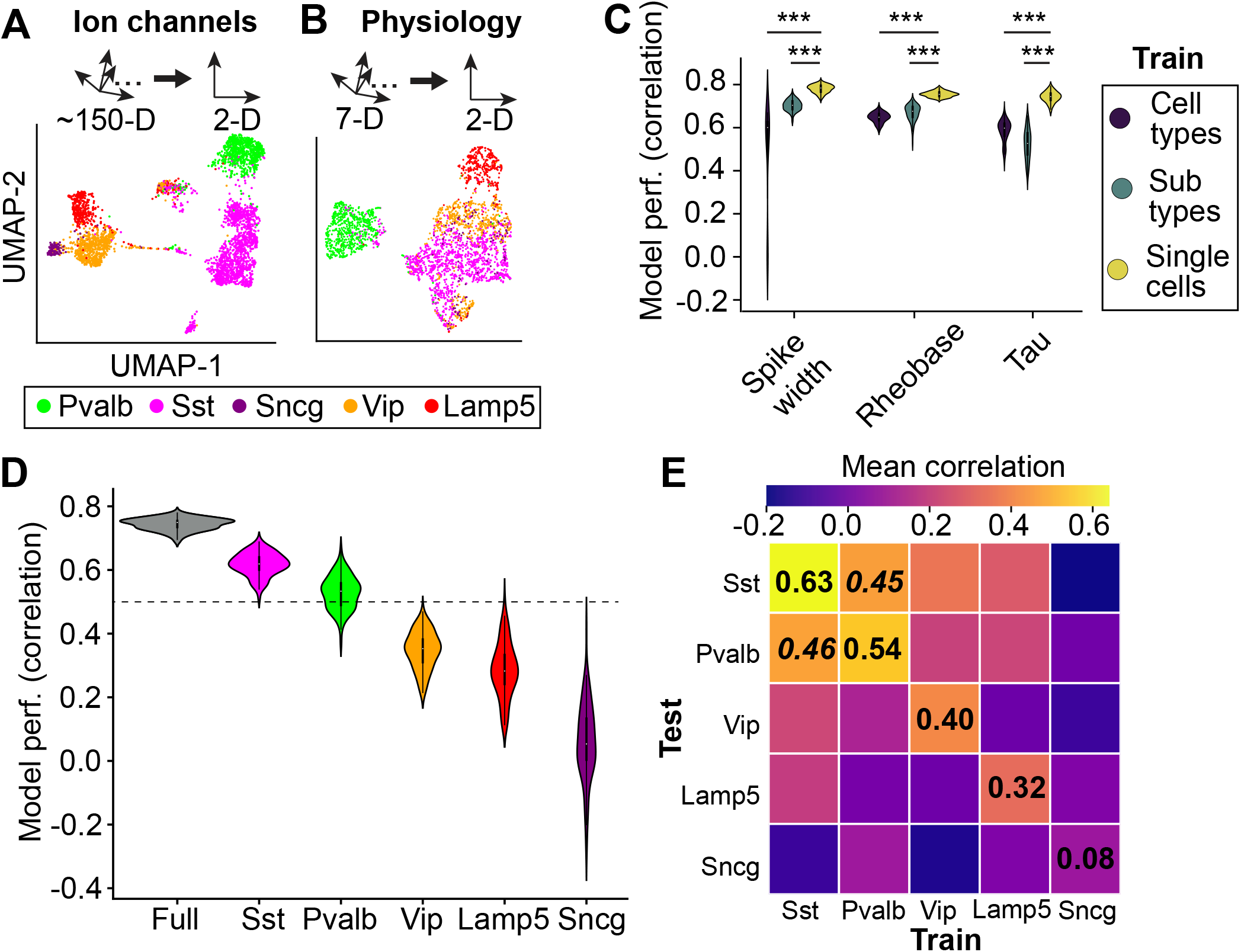
Single-cell gene expression improves prediction of physiology relative to cell type averaging and enables within-cell-type physiology prediction. A. Ion channel gene expression (*n* = 3651 cells) and **B.** Physiology (*n* = 3268 cells) delineates canonical cell types in dimensionally-reduced UMAP space. **C.** Discretizing single cell datasets into sub-type or cell type averages reduces model accuracy on held-out single cell data for all physiology features we examined, a subset of which are shown here. ***: *p* < 0.001, unpaired two-sided *t*-test with Bonferroni correction. **D.** Within-cell-type model predictive accuracy (spike width). Spike width can be predicted within Pvalb and Sst interneuron populations with mean correlation of *R* > 0.5 between model predictions versus actual spike width (*n* = 100 train-test splits). **E.** Spike width performance within and across cell types. The total number of cells per cell type with data for each physiology feature – before dividing into 80/20 train-test splits – is given in Supplemental Table S1. Confidence intervals for within-cell-type analyses as determined from 100 train test splits in order from Sst to Sncg: [0.57,0.70], [0.47,0.63], [0.29,0.50], [0.16,0.47], and [- 0.17,0.35]. For cross-cell-type analyses, confidence intervals were determined by 100 bootstrap samples of the training dataset. Confidence intervals for: training on Pvalb and testing on Sst: [0.31,0.48]; training on Sst and testing on Pvalb: [0.39,0.47].

We trained models on cell type averages at the cell-type (e.g., Sst) and subtype (e.g., Sst Tac1 Tacr3) levels and evaluated models on held-out single cell data. Models trained on single cells outperformed those trained on cell type averages at both the cell-type and subtype level for all physiology features we examined (Figure 2C; Supplemental Figures S3A and S3B). Thus, single-cell ion channel gene expression contains signal predictive of physiology that is not captured by cell type averages.

Nevertheless, model predictions based on cell-type- or subtype-averaged gene expression were well above *R* > 0.5 for several physiology features, suggesting that cell type differences contributed significantly to model predictions. To explicitly test this, we trained models to predict physiology from a categorical predictor alone that denoted which cell type or subtype a neuron belonged to (‘one-hot’ encoding). In line with our previous results, physiology was well-predicted from cell type or subtype identity alone (Supplemental Figures S3C and S3D).

We added ion channel gene expression to our categorical model to test if this added predictive information not captured by cell type identity, finding that this significantly improved model performance (Supplemental Figure S3C). Confoundingly, we found that the addition of ion channel gene expression to cell subtype information did not improve model predictions, and sometimes even marginally reduced model performance (Supplemental Figure S3D). This contrasted with our previous finding that models trained on cell subtype averages performed worse than those trained on single cells. We speculate that this difference arose from the greater reliability of information content in cell subtype assignments, which are derived from mapping whole genomes to a reference taxonomy of thousands of cells,^16,49^ relative to ion channel gene expression obtained from single cells.

In general, cell-type- and subtype-averaged models were highly sparse. For example, canonical channels thought to underlie fast-spiking of Pvalb interneurons—encoded by *KCNC1, KCNC3*, *SCN1A*, and *SCN8A—*were not selected by cell-type- or subtype-averaged models as related to spike width, but were selected by single-cell models (Supplemental Table S4). This finding is further underscored by a report that models based on cell-type-averaged data did not identify *KCNC1* as related to spike width.^17^ By contrast, adding cell type or subtype as a ‘one-hot’ categorical predictor to single-cell ion channel gene expression resulted in more non-zero model coefficients (Supplemental Table S4). For example, controlling for cell type or subtype unmasked negative correlations between *HCN1* expression and spike width, consistent with reports that axon-specific expression of HCN1 in fast-spiking interneurons facilitates faster spiking.^50^

To further probe the extent to which our model could parse covariation between ion channel gene expression and physiology within-cell-type, we next tested if we could predict physiology from gene expression within a given cell type. We observed that within-cell-type variation in gene expression significantly predicted at least one physiology feature for each cell type (Supplemental Figure S4).

As in the full dataset, spike width was best predicted within-cell-type, particularly within Sst and Pvalb interneuron populations (Figure 2D). Genes associated with narrower spike widths in both Sst and Pvalb interneurons encoded canonical potassium and sodium channels that facilitate high frequency firing: Kv3.1 (*KCNC1*), Kv3.3 (*KCNC3*), and Nav1.1 (*SCN1A*) and Nav1.6 (*SCN8A*) (Supplemental Figure S5; Supplemental Table S5). These channels are typically associated with fast-spiking in Pvalb interneurons (but see: Nav1.1 deletion in Sst interneurons impairs excitability^51^) and are often used as cell type marker genes for Pvalb neurons.^40,42^ Our finding that graded expression of these channels predicted spike width within Sst interneurons therefore suggests that the molecular regulation of interneuron spike width may be more general than previously appreciated.

We asked if similarities in gene expression-spike width relationships were strong enough to predict across cell types. We found that models generalized well across Pvalb and Sst interneurons, but not to other cell types (Figure 2E).

By contrast, tau was predicted well within Sst interneurons, but not within Pvalb interneurons (Supplemental Figure S4). Combined with the failure of spike width models to generalize well across cell types other than Pvalb and Sst interneurons, our findings illustrate that gene-physiology relationships are heterogeneous across cell types and physiology features.

### Synaptic and GPCR gene sets are co-regulated with ion channel genes and predict physiology with equivalent model accuracy

We next asked if ion channel genes were uniquely predictive of physiology relative to other gene sets (Figure 3A). To address this, we selected two alternative gene sets based on the criteria that each set contained a comparable number of members and a similar capacity to delineate cell type: synaptic markers and G-protein coupled receptors (GPCRs) (Supplemental Figure S6).^9,52^ We hypothesized that the expression of synaptic or GPCR genes may be sufficiently coupled to ion channel gene expression to reliably predict physiological variation.

**Figure 3.**
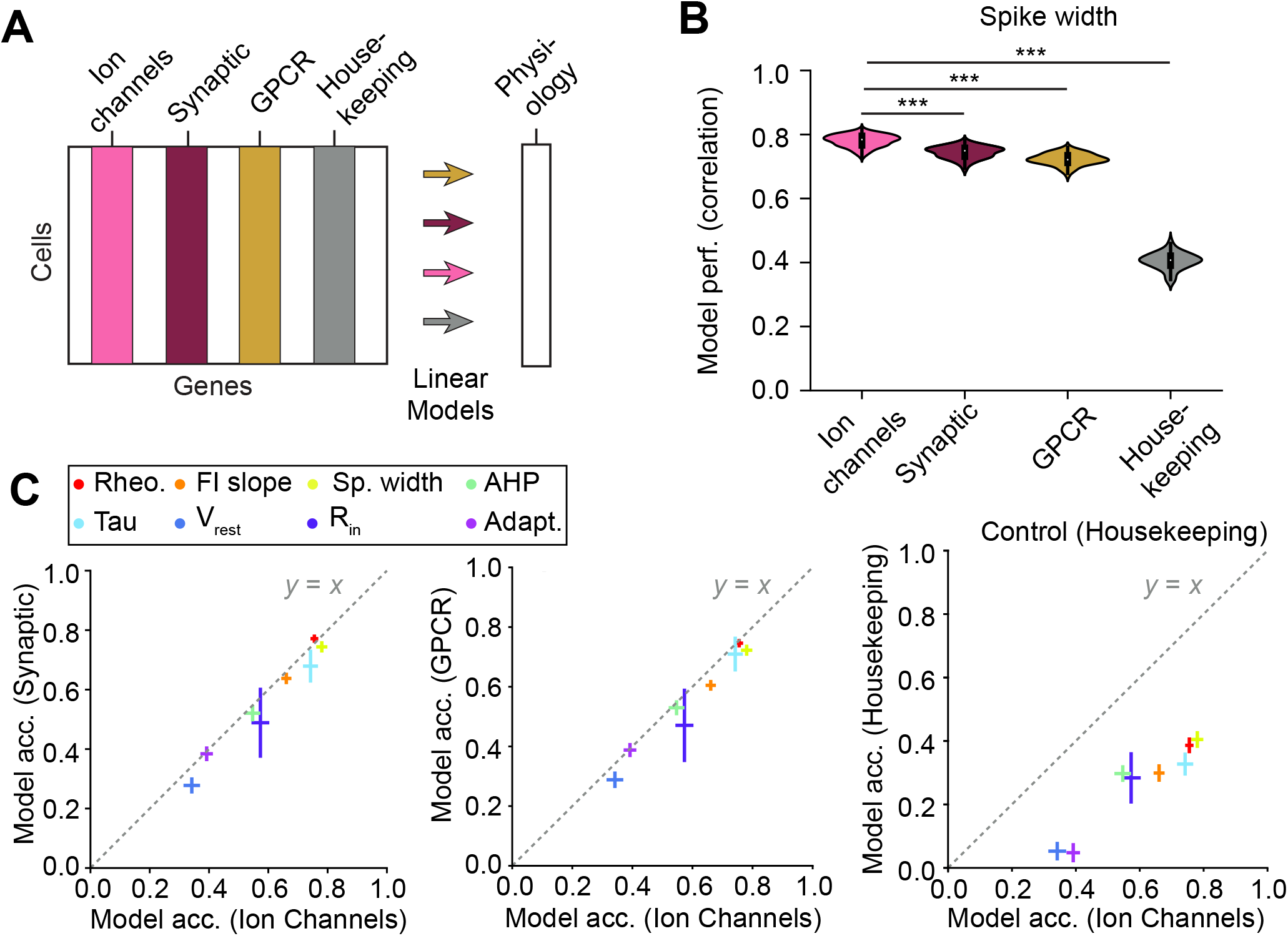
Synaptic and GPCR gene sets also predict variation in physiology. **A**. Linear models were fit to predict physiology from the expression of gene sets: ion channels, synaptic, and GPCR. **B.** Synaptic and GPCR gene sets predict spike width numerically comparably to ion channels, with statistically higher predictive accuracy of ion channel gene expression. Synaptic, GPCR, and ion channels all predict spike width significantly better than housekeeping genes. *** denotes *p* < 0.001; two-sided, paired *t*-test with Bonferroni correction. **C.** Synaptic and **D.** GPCR gene sets predict physiology numerically comparably to ion channels across all considered physiology features. However, ion channels statistically outperformed synaptic and GPCR genes for prediction of all physiology features except for rheobase (all *p* < 0.001; two-sided, paired *t*-test with Bonferroni correction) with two exceptions: Synaptic gene expression significantly better predicted Rheobase (*p* < 0.001) and did not significantly differ in prediction of Adaptation (*p* = 0.14) relative to ion channel genes. Ion channels predictive accuracy was much higher than a control housekeeping set (right; all *p* < 0.001).

Indeed, models trained on the expression of synaptic genes or GPCR genes performed comparably to models trained on ion channel gene expression for every physiology feature we examined (Figure 3B and 3C left, middle). Critically, prediction from ion channel, synaptic, and GPCR gene sets all outperformed a gene set with a similar number of members but that would not be reasonably expected to relate to physiology, housekeeping genes (Figure 3B and 3C, right). Moreover, within-cell-type variation in GPCR and synaptic gene expression each successfully predicted physiology for some combinations of cell types and physiology features (Supplemental Figure S4).

We clustered GPCR and synaptic genes based on assigned coefficients across predictive models. This revealed a ‘fast-spiking’ module of synaptic genes whose expression was associated with narrower action potentials and steeper FI slopes (Supplemental Table S2). Expression of genes encoding synaptic proteins integral to fast vesicle release—*SYT2* and *VAMP1*—were revealed to negatively correlate to spike width by models fit to the full dataset and to Pvalb interneurons alone (Supplemental Tables S2 and S5). This adds a functional dimension to reports that *SYT2* and *VAMP1* expression correlates with fast-spiking Kv3 channel expression^38^ and that *SYT2* up-regulation coincides with the development of fast-spiking in Pvalb interneurons.^39^

For GPCR-physiology models, the gene encoding mGluR1 (*GRM1*) was identified as related to rheobase by models fit to the full dataset and to cell types alone. Consistent with this, 1) mGlur1 has been previously associated with homeostatic changes in intrinsic physiology through interactions with HCN channels ^53,54^ and 2) knockout of *GRM1* decreases input resistance,^55^ in line with the negative correlation to rheobase identified by GPCR-physiology models (Supplemental Tables S2 and S5).

Therefore, our results indicate that synaptic and GPCR gene expression covaries with intrinsic physiology. Comparison of our modeling results with prior research suggests that this covariation arises from coordinated regulation of synaptic and intrinsic physiology, and may even indicate direct functional interactions between ion channels and other proteins that shape neuronal responses to synaptic input like GPCRs.

### Information for prediction of physiology from GPCR and synaptic gene sets is fully contained in ion channel gene expression

We next asked if synaptic or GPCR genes contained different information than ion channel genes for prediction of physiology. We did this by residualizing GPCR and synaptic gene expression for ion channel gene expression (Figures 4A and 4B). We found that ion channel gene expression explained significant variance in synaptic and GPCR gene expression when compared to control housekeeping genes (Supplemental Figure S7). Moreover, residualizing synaptic and GPCR gene expression data for ion channel gene expression completely abolished model performance (Figure 4C).

**Figure 4.**
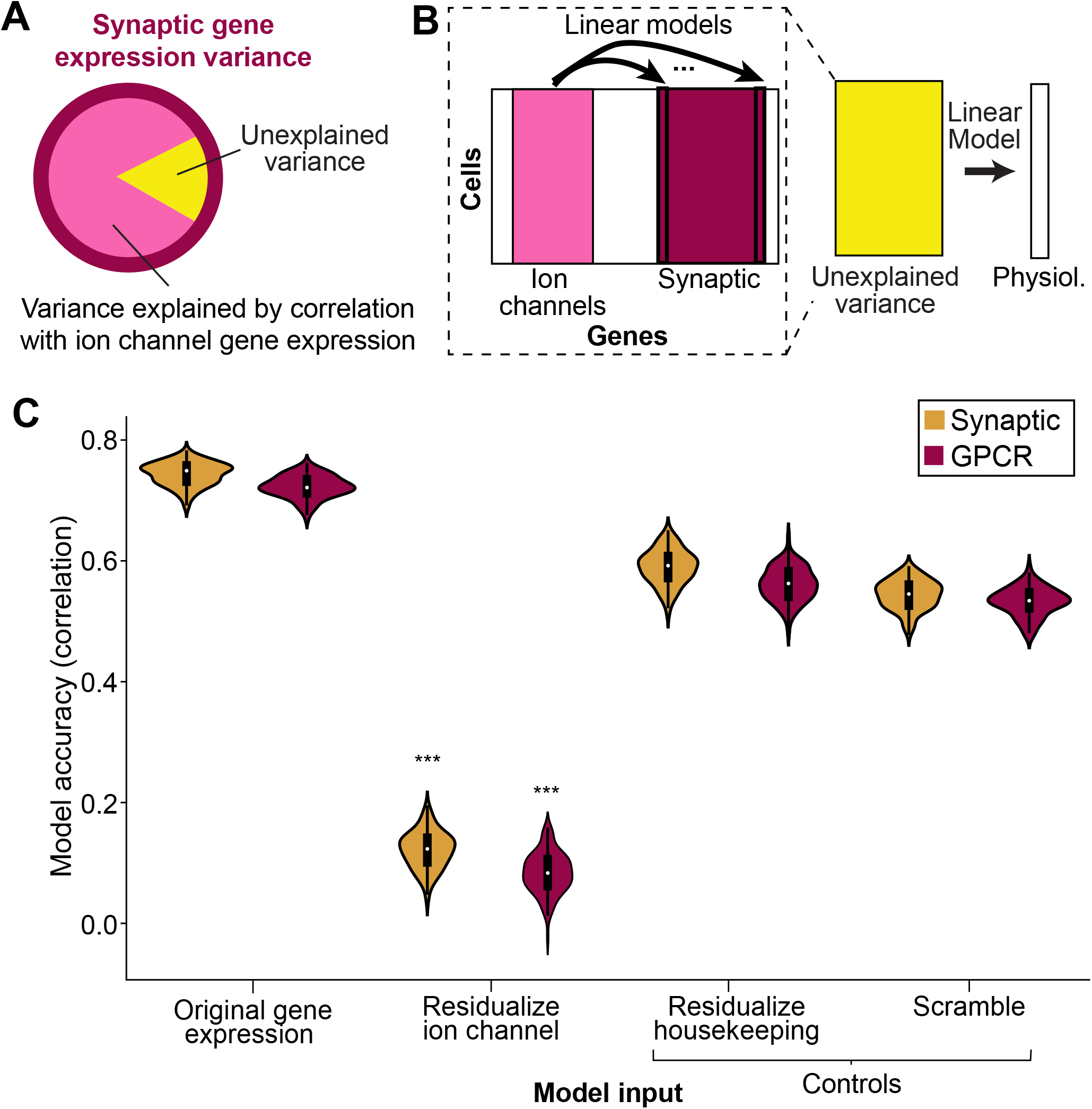
Synaptic and GPCR gene sets are co-regulated with ion channel genes and predict physiology with equivalent model accuracy. A. Concept: Some subset of variation in synaptic (or GPCR) gene expression covaries with ion channel gene expression, while some variation cannot be explained by ion channel gene expression. If synaptic (or GPCR) gene expression uniquely explains variation in physiology that is not captured by ion channel gene expression, this residualized synaptic or GPCR gene expression is still predictive of physiology. **B.** Implementation: Linear models were fit to predict the expression of synaptic (and GPCR) genes from ion channel gene expression. Unexplained variance that could not be fit by ion channel gene expression was fit to a linear model predicting physiology. **C.** Model predictive power was abolished after subtracting variance in synaptic and GPCR gene expression explained by ion channel gene expression. Subtracting variance explained by housekeeping, or scrambling model fits from ion channel gene expression, did not abolish model predictive power to the same degree. *** denotes *p* < 0.001, two-sided unpaired *t*-test for all pairwise comparisons within a given gene set.

It is possible that residualizing GPCR and synaptic gene expression for ion channel gene expression abolished model performance by making gene expression data noisier. To test this potential limitation, we subtracted off 1) model predictions derived from housekeeping gene expression and 2) scrambled model predictions such that they were misaligned with gene expression data. When we fit physiology data to this noisy gene expression data, in both cases, this only slightly reduced model performance (Figure 4C). This demonstrates that residualizing GPCR and synaptic gene expression for ion channel gene expression does not abolish model performance simply by introducing noise, but instead by eliminating the explanatory power from GPCR and synaptic gene expression data.

Thus, we conclude that while ion channel gene expression likely relates most directly to physiology *a priori*, co-regulated synaptic and GPCR gene sets closely track physiological function.

### Ion channel gene expression predicts physiology within human cortical pyramidal neurons

We next tested if this approach could be extended to predict physiology from gene expression in a separate, human pyramidal neuron Patch-Seq dataset.^30^ This dataset has unique constraints that tested the robustness of our approach: 1) it is an order of magnitude smaller than the mouse dataset (∼300 human versus ∼3000 mouse neurons); and 2) it is comprised of Layer 2/3 (L2/3) cortical pyramidal cells, which are more molecularly and physiologically homogeneous than cortical interneurons (Figure 5A).^56^

**Figure 5.**
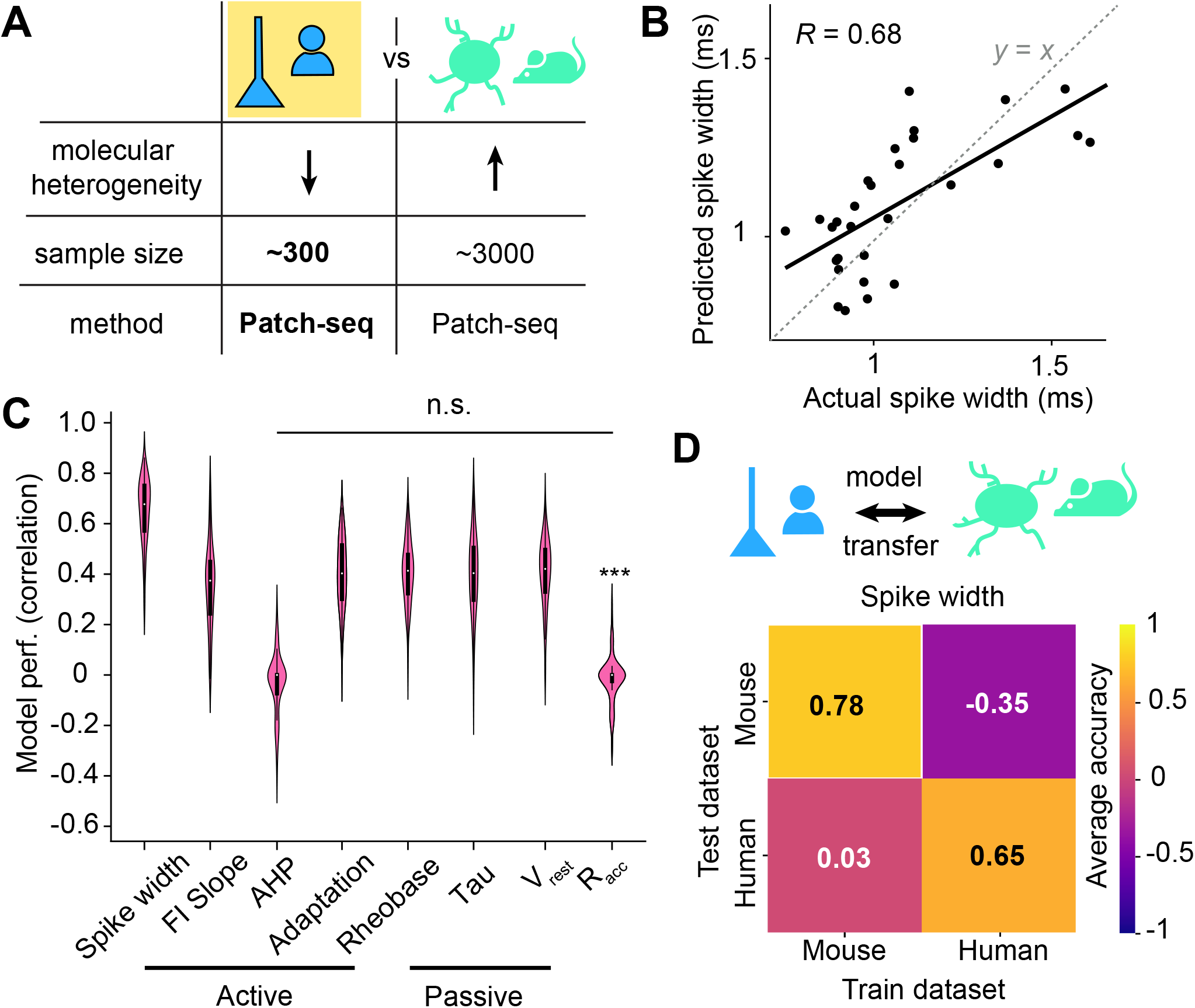
Ion channel gene expression predicts physiology within human pyramidal neurons, with distinctive gene-physiology relationships from mouse interneurons. A. Dataset overview: Dataset is comprised of human pyramidal neurons. Key factors that might influence model predictive accuracy in this dataset include reduced sample size (∼300 human neurons versus ∼3000 mouse neurons) and reduced molecular heterogeneity of pyramidal neurons versus interneurons. B. Ion channel gene expression predicts spike width of human cortical pyramidal neurons (Pearson’s *R* = 0.68; *n* = 58 cells). Dashed line denotes *y*=*x* and displayed data comprises held-out test samples. C. Spike width is predicted most accurately relative to other physiology features (mean *R* = 0.65), followed by input resistance (mean *R* = 0.50) and rheobase (mean *R* = 0.40), all *n* = 100 train-test splits. All physiology features were significantly better predicted than the control measurement, Racc (all *p* < 0.001), with the exception of AHP (*p* = 1.84) on a two-sided, unpaired *t*-test with Bonferroni correction. D. Model transfer across datasets. While spike width was well-predicted within both Patch-Seq datasets, models trained on one dataset did not generalize to the other dataset. 95% confidence intervals (*n* = 100 train-test splits): mouse [0.74,0.80] and human [0.32,0.82]. For cross-dataset analyses, confidence intervals were determined by *n* = 100 bootstrap samples of the training dataset: training on mouse and testing on human [-0.06,-0.16] and training on human and testing on mouse [-0.42,0.11].

Despite these constraints, a model trained on human cortical pyramidal neurons could accurately predict the spike width of held-out samples (Figure 5B). As in the mouse interneuron dataset, spike width was the best-predicted physiology feature in human pyramidal neurons and the only feature that met our criteria (*R* > 0.5) for further examination of model coefficients (Figure 5C).

Genes selected by our human pyramidal neuron spike width model were biologically plausible (Supplemental Table S6). The expression of Kv4 channels (*KCND2* and *KCND3*) and of an electronically silent subunit of Kv2 channels, Kv6.3 (*KCNG3*) each was strongly negatively correlated with spike width, in line with previous interventional work in mouse pyramidal neurons.^57–60^

By contrast, the expression of Kv4 channels was associated with wider spikes in mouse interneurons (Supplemental Table S2). Correlational work in Pvalb interneurons aligns with these mouse interneuron modeling results, demonstrating that downregulation of Kv4 channels coincides with the development of fast-spiking abilities; however, to our knowledge, this has not yet been mechanistically explored.^39^

In general, gene expression-physiology relationships were distinct between L2/3 human pyramidal neurons and mouse interneurons (Supplemental Table S7; Supplemental Figure S8), and model performance did not transfer between datasets (Figure 5D). In fact, model predictions of mouse interneuron spike width were anti-correlated with measured spike width after training on human pyramidal neurons; this is likely due to opposing relationships between Kv4 channel expression and spike width across the two datasets.

The only conserved gene-physiology relationship in both human pyramidal neuron and mouse interneuron spike width models involved *SCN1A* (Nav1.1) expression. This was remarkable because, in mouse L2/3, Nav1.1 channels are primarily restricted to fast-spiking interneurons^42^ (see also^61^), and knockout of *SCN1A* does not affect excitability in mouse pyramidal neurons.^51^ Our model revealed that human pyramidal neurons express *SCN1A*, and that this expression negatively correlates to spike width, as in mouse fast-spiking interneurons. One previous study demonstrated at the protein level that human cortical pyramidal neurons express Nav1.1,^62^ but a later study could not replicate this finding.^63^ Here, our results extend this work to suggest the functional relevance of Nav1.1 channels for determining spike width within L2/3 human pyramidal neurons.

## Discussion

Single-cell transcriptomics promises unprecedented insight into the molecular underpinnings of biological systems. However, the extent to which single-cell transcriptomic data can be leveraged to propose hypotheses about gene-function relationships in a heterogeneous population of cells is unknown. Even in the rare case where transcriptomic data and functional measurements can be obtained from the same cell (e.g., Patch-Seq), technical noise can make it challenging to connect these two modalities. Therefore, previous approaches to analyzing Patch-Seq data have primarily relied on working with cell type averages to reduce noise. We asked if we could connect transcriptomics to physiology in single cells, to provide a more detailed understanding of cellular relationships.

### Single-cell transcriptomic data predicts single-cell physiology

We use linear modeling to show in both mouse interneurons and human pyramidal neurons that transcriptomically-measured ion channel gene expression can predict variation in physiology at single-cell resolution (Figures 1 and 5). Critically, we find that single-cell gene expression predicts single-cell physiology better than cell-type-averaged gene expression data (Figure 2). This suggests that previous approaches of de-noising single-cell gene expression data by averaging on cell types^9,16–19^ sacrifice signal that is predictive of biological function.

### Model validation and comparison to previous research

One other study by Cadwell and colleagues^8^ applied linear models to Patch-Seq data to predict physiology at single-cell resolution. As this study was one of the first to establish the Patch-Seq method (see also: Földy et al.^9^ and Fuzik et al.^10^), sample sizes were orders of magnitude smaller than the more recent publicly-available Patch-Seq datasets.^4,16^ Remarkably, the authors still found that single-cell gene expression can predict some physiology features, although spike width and tau could not be predicted.

By contrast, our models predicted these physiology features with some of the highest accuracy within both mouse interneurons and human pyramidal neurons. As the model in Cadwell et al. was pooled on interneurons and pyramidal neurons, we speculate that failure to build a predictive model of spike width may be due to differential molecular regulation of this feature by these cell classes.^18^ Indeed, we found that spike width was best predicted within both interneurons and pyramidal neurons (Figures 1 and 5), but nearly all identified gene expression- physiology relationships were distinct (Supplemental Table S7; Supplemental Figure S8; but see also discussion of Nav1.1 below), and models did not generalize across cell classes (Figure 5). Because the datasets that we analyzed also varied by species, we cannot rule out species differences as a primary driver of heterogeneity. However, even within mouse interneurons, cell types had heterogeneous gene expression-physiology relationships: tau varied continuously within the Sst cell type in a manner that could be predicted from gene expression data, but was remarkably homogeneous within Pvalb interneurons. This highlights that with enough data, single-cell heterogeneity (particularly within cell types), can be an avenue for exploration rather than a potential point of modeling failure.

Another key difference between our model and that of Cadwell et al. was the gene set used. Cadwell et al. built their model on the most differentially expressed genes, none of which were ion channel genes. By contrast, we placed a ‘biological prior’ on our model by restricting it to ion channel genes only. The fact that we were able to build generalizable models predicting physiology features from single-cell ion channel gene expression is nontrivial, given the profound degeneracy in neuronal ion channel expression described by previous studies.^11,33,64^ These seminal studies showed that multiple different combinations of ion channels can produce the same physiological output. This might be expected to preclude identifying general ion channel-physiology relationships across many different neurons. Moreover, our model identified biologically plausible examples where the expression of two different ion channels were equivalently related to one physiology feature, but differently related to another physiology feature (e.g., BK/Kv3 expression: shared relationship to spike width, only BK related to AHP; Kv1.1-2/DPP6 expression: shared relationship to spike width, opposite relationship to rheobase). Thus, our modeling results add to a growing body of evidence that ion channel degeneracy is not simply redundancy, but instead represents functional overlap in some contributions to physiology.^33,34^

To further validate our model, we probed if identified gene expression-physiology relationships were biologically plausible by comparing our findings to previous studies leveraging genetic knockouts, pharmacology, and RT-PCR. We found that gene detection with transcriptomics is sufficient to replicate several specific ion channel-physiology relationships. Our mouse interneuron model recapitulated correlations between: 1) Kv3 and Nav1.1 channels and fast-spiking^6,37–43^, 2) Kv1.1-2 channels and higher rheobase and narrower spikes^18,42^, and 3) between HCN1 and a faster tau.^6^ Our human pyramidal neuron model recapitulated findings from mouse cortical pyramidal neurons that overexpression of Kv6.3 or Kv4.2 narrows spike widths.^57–60^

### Utility of transcriptomics for functional insight over previous methods

What can be gained by using transcriptomics to better understand function, and specifically physiology? At a technical level, transcriptomics is a far less laborious technique than pharmacology or RT-PCR. This is especially critical for maximizing data collection from rare samples, such as human tissue.^30^

Moreover, access to information about the whole genome enables a multi-dimensional view of the molecular regulation of physiology. This perspective is especially useful in light of growing evidence of coordination between ion channels, GPCRs, and synaptic proteins to serve normal physiological function. For example, 1) GPCR-mediated neuromodulatory input is indispensable for adjusting ion channel expression to maintain robust computational power in the context of fluctuating network activity levels;^65–67^ 2) Disruption of direct interactions between ion channel and synaptic or GPCR proteins leads to aberrant physiology and, ultimately, to disease;^68^ 3) Physiologically-distinctive cell types express different repertoires of synaptic proteins^9,52^ and 4) ion channel and synaptic gene expression covary.^38^

Here, we extended this work by demonstrating that synaptic and GPCR gene expression is sufficiently coupled to ion channel gene expression to enable accurate prediction of physiology from these gene sets (Figure 3). Our correlational analysis cannot rule out the possibility that GPCR and synaptic proteins derive their predictive power from direct causal relationships to physiology. However, decades of work indicating that ion channels are the key determinants of intrinsic physiology, combined with our findings that GPCR and synaptic gene sets did not contain information predictive of physiology beyond that already contained in ion channel genes (Figure 4), leads us to conclude otherwise.

### Novel ion channel-physiology relationships

Our linear modeling approach revealed multiple novel gene expression-physiology relationships that can be tested in subsequent studies. Within mouse interneurons, we found that several ion channels that were previously thought to be primarily restricted to fast-spiking interneurons^42,69^ were also expressed within non-fast-spiking Sst interneurons. Expression of each of these genes, which encoded channels for Kv3.1-3.3, Nav1.1, and Nav1.6, was negatively correlated to spike width, as in fast-spiking interneurons. Notably, these genes were identified as related to spike width only by models based on single-cell gene expression data, and not by models fit to cell-type averages. This suggests that covariance between the expression of fast-spiking ion channel genes and spike width has a prominent within-cell-type component and underscores the insight gained from retaining single-cell resolution.

We also found that expression of the gene encoding Nav1.1 negatively correlated to spike width within L2/3 human pyramidal neurons. This points towards a previously disputed cross-species difference, since previous work in L2/3 of mouse neocortex has found that Nav1.1 ion channels are almost exclusively expressed by fast-spiking interneurons.^42^ Surprisingly, one previous study found that Nav1.1 was expressed by L2/3 human pyramidal neurons using protein immunolabeling,^62^ but this potential cross-species difference failed to replicate in a subsequent study.^63^ In addition, potential functional impacts of any Nav1.1 expression on human pyramidal neuron physiology has not yet been explored to our knowledge.

Our finding that Nav1.1 expression is associated with narrower spike widths in L2/3 human pyramidal neurons in a manner analogous to fast-spiking interneurons suggests that Nav1.1 channels could contribute to the faster action potential kinetics of human relative to mouse pyramidal neurons.^70^ In fact, faster action potential kinetics, including narrower spike widths, has been proposed to facilitate higher-order cognition and faster processing speeds of humans and non-human primates relative to other mammalian species like mice.^70,71^ However, future interventional work directly comparing human and mouse cortical pyramidal neurons will be necessary to test if Nav1.1 expression could be a potential molecular substrate of these cross-species physiological differences.

### Application to understanding neuronal function in disease

Our models identify genes that robustly predict physiological variation in well-powered datasets spanning multiple species and cell types. Thus, future work could apply our results to interpret smaller, specialized datasets, such as those focused on disease mechanisms. Knockout of a disease risk gene is likely to trigger many transcriptomic changes, a subset of which are functionally relevant. Our model could be used as a guide to determine which transcriptomic changes likely underly disease pathophysiology. In particular, impaired cortical neuron excitability is associated with epilepsy,^72^ autism,^73^ schizophrenia,^74^ obsessive-compulsive disorder,^75^ and other disorders,^76,77^ and our model could be used to identify potential molecular contributors to these disease phenotypes.

## Methods

### Patch-Seq datasets

We worked with Patch-Seq datasets collected by the Allen Institute that have been previously analyzed in two different studies: Gouwens et al. 2020^16^ (mouse interneurons across the cortical layers) and Berg et al. 2021^4^ (human pyramidal neurons in L2/3). The mouse interneuron Patch-Seq dataset comprised 3,855 cortical GABAergic interneurons that passed QC criterion and consistently mapped to a transcriptomic type and thus were included in the released counts matrix. The human pyramidal neuron Patch-Seq dataset was collected from live, neurosurgically-resected human tissue and comprised 385 neurons that passed QC.

### Physiology features

We used precomputed physiology features when available.^16^ Otherwise, we calculated features using the Allen Institute Intrinsic Physiology Feature Extractor (IPFX) Python package. We created a physiology dataset using `ipfx.dataset.create.create_ephy_dataset`, followed by quality control (`ipfx.utilities.drop_failed_sweeps`) and filtering on long square (1s stimulus) sweeps (`data_set.filtered_sweep_table(stimuli=data_set.ontology.long_square_na mes`)). We used `ipfx.stimulus_protocol_analysis.LongSquareAnalysis.analyze()` to extract physiology features. Not all neurons had quality data available for every physiology feature we examined. Generally, spiking features had a reduced sample size, as some neurons did not meet the minimum threshold of elicited spikes needed to analyze spiking features on any trace (see detailed criteria below). See Supplemental Table S1 for sample sizes for each physiology feature.

For details on physiology, see: Allen Cell Types Database Technical White Paper: Electrophysiology (https://celltypes.brain-map.org/) and previous Patch-Seq papers’ Methods.^4,16^ Briefly, all physiology features were calculated from long-square pulses that had a duration of 1 second. Spike width was calculated from the first trace that elicited five spikes, defined as the duration of time between half-height of the rising versus falling phases of the action potential, and averaged from all spikes in the trace. Rheobase was defined as the minimum amount of current injected that elicited at least one spike. Membrane time constant (tau) was calculated by fitting a decaying exponential to the return to resting membrane potential after a hyperpolarizing voltage step. FI slope was calculated by regressing the spike frequency from the amount of current injected. AHP was defined as the most hyperpolarized value of the membrane potential in the 5 ms time window after the action potential peak. Adaption was defined as the average fractional change in duration of subsequent ISIs across the spike train. Resting membrane potential and access resistance were determined soon after break-in and before the first stimulus injections. A full matrix of physiology features and values across cells is given in Supplemental Table S8.

### Transcriptomic data preprocessing

We first subset the transcriptomic count matrices on genes that were expressed at non-zero levels in at least 1% of cells in the dataset. To account for different sequencing depths, we within-cell-log-normalized the raw gene expression counts data for genes that remained after the previous filtering step. As the logarithm function goes to negative infinity at 0, we added 1 to every entry of our raw gene expression counts matrix before log normalizing, making the final value for each gene in a given cell: log(counts+1 / total CPM of RNA in that cell). In addition to the technical advantages of within-cell-normalization, relative values of ion channel levels are an important determinant of physiology,^11,37^ a feature that would not be captured by normalizing across cells.

### Cell type definitions

For our cell type analyses on the mouse interneuron dataset, we used assignments provided in the publicly available metadata from the Allen Institute (https://portal.brain-map.org/cell-types/classes/multimodal-characterization). For further details on how these cell types were assigned, see the original publication: Gouwens et al. 2020.^16^ For cell type assignments, see Supplemental Table S8.

In each dataset, there were granular and coarse cell type definitions, which at times conflicted. To resolve these conflicts, we based our analyses on the granular cell type definition, and assigned cell type based on the first gene listed in the definition. That is, a cell labeled as Sst Tac2 Myh4 would be classified as Sst.

Two cell subtypes, both labeled principally by Serpinf1, were spread across both Vip and Sncg subtypes with no apparent pattern. As this only comprised *n* = 40 cells in the full dataset of *n* = 3691 cells and could not be unambiguously assigned to an overarching cell type, we did not analyze these cells separately for our cell type restricted analysis. Instead, we only considered types principally marked by Sst, Pvalb, Vip, Lamp5, and Sncg, as in Gouwens et al.^16^

### Gene sets

Housekeeping genes were selected from the top ∼150 most stably expressed genes previously determined from 11 publicly available scRNA-seq datasets spanning human and mouse, cell types, and tissues (including neural tissue).^78^ Synaptic, GPCR, and ion channel genes were obtained from HUGO (https://www.genenames.org/), an online repository that annotates and groups genes based on function. Ion channel genes were manually curated from HUGO to add auxiliary subunits that are key to function and expression of ion channel genes.

Final gene lists for each dataset and gene set, after restricting to genes that were expressed at non-zero levels in at least 1% of cells, are provided in Supplemental Table S9.

### L1-regularized linear model

We fit L1-regularized linear models using `sklearn.linear_model.LassoCV`, allowing for an intercept (`fit_intercept = True`). In the outer loop, we subsetted our dataset into 100 different train-test splits to determine if well-fit models were largely invariant to the particular train-test split of the data and thus robust. We chose this number of train-test splits based on empirical testing (Supplemental Figure S1). In the mouse interneuron dataset, we divided data into 80% train and 20% test. To allow for fitting on more training data in the smaller human pyramidal neuron dataset, we split the data into 90% train and 10% test.

In general, in the inner loop we used 10-fold cross-validation on the training data to pick the regularization strength (alpha) for L1-penalized, or LASSO, regression.^31^ In the case of fitting our model on broad cell type averages in our interneuron dataset, of which there are only 5 (Pvalb, Sst, Sncg, Vip, and Lamp5), we used 3-fold cross-validation because the data could not be cross-validated on 10-folds. However, when we trained our model on cell subtype averages (e.g., Sst Tac1 Tacr3), we again used 10-fold cross-validation. Because we still found reduced model performance in this context, we conclude that the reduction in model performance we obtained from training on cell type averages was not due to a reduced number of cross-validation folds.

LASSO regression enforces sparsity of model fits, such that high-magnitude coefficients are penalized and certain coefficients are dropped to zero. This mitigates overfitting concerns and reduces the set of features to a set that are ‘selected’ as useful to the model, improving model interpretability.

### L2-regularized and unregularized models

To empirically confirm the validity of our LASSO regression approach, we also fit models using L2 regularization (i.e., Ridge^32^) and without regularization via least-squares. To fit Ridge models, we used `sklearn.linear_model.RidgeCV` with (`fit_intercept = True`). As with our LASSO models, we applied a nested cross-validation procedure with 10-fold cros- validation to choose regularization strength, followed by an outer loop where we divided data into 100 train-test splits. For least-squares linear regression, we used `sklearn.linear_model` with (`fit_intercept = True`). Again, we fit linear models to 100 train-test splits of the data.

### Model coefficient interpretation and visualization

To identify modules of genes that displayed similar relationships to physiology features, we clustered genes based on their assigned coefficients across physiology models. To ensure the robustness of our results, we took several measures before clustering. First, we only analyzed model coefficients for gene-physiology models that were reasonably predictive, meaning measured versus predicted physiology had mean Pearson’s *R* > 0.5 across train-test splits. Second, within a given gene-physiology model, we only included genes that showed largely consistent relationships to physiology across train-test splits. Specifically, we only included genes that were assigned non-zero, same-signed coefficients on 80% of train-test splits. Finally, to obtain smoothed estimates of model coefficients, we averaged assigned coefficients across 100 train-test splits for genes that met the 80% criterion.

We performed Ward’s hierarchical clustering on the resulting gene × physiology matrix using the clustermap package in the `seaborn` library^79^ in Python. We cut the tree to yield 4 clusters using `scipy.cluster.hierarchy.cut_tree` on our `clustermap`-generated dendrogram.

Several of our analyses involved comparing model coefficients across cell types and physiology features. In both cases, physiology can vary in magnitude both as a function of different physiology features that have different measurement units (e.g., rheobase versus resting potential) or as a function of cell types that have appreciable differences in mean physiology values (e.g., Pvalb interneurons have 2-fold narrower spike widths than most interneurons in the dataset on average). Thus, to correct for differences in the magnitude of the output variable across cell types or physiology features while still maintaining the direction of the relationship between a given gene and physiology (i.e., sign of the coefficient), we normalized coefficients to the absolute maximum within a cell type or physiology feature. We then visualized absolute maximum-scaled model coefficients using the `clustermap` package, plotting genes on the *y*-axis and cell types or physiology features on the *x*-axis depending on the analysis.

### Visualization of high-dimensional single-cell gene expression and physiology data

To visualize high-dimensional single-cell gene expression data or physiology data (i.e., for multiple physiology features), we employed dimensionality reduction. For gene expression, we log-normalized transcriptomic data on the subset of genes of interest (e.g., ion channels) and reduced to a two-dimensional embedding using the `umap-learn` package^80^ in Python. For physiology features, we built a cell × physiology feature matrix, only including cells that had values for all physiology features in the matrix. We included all examined physiology features in the matrix: rheobase, spike width, tau, FI slope, AHP, Adaptation, and Vrest. To account for different measurement units, we z-scored within a given physiology feature across cells, and then dimensionally reduced the resultant matrix via `umap-learn`.

### One hot-encoding of cell type

To account for variance explained by cell type, we encoded cell type or subtype as a categorical variable using one-hot encoding. That is, we appended a column for each cell type or subtype to our input matrix of gene expression data. Each cell was assigned a 1 to the entry corresponding to its cell type or subtype (see “Cell type definitions” above). To control for differences in magnitude between log-normalized gene expression and one-hot encoding columns, we scaled each column across cells such that each input feature had unit variance and mean 0. To analyze ion channel-gene expression relationships after accounting for cell type, we excluded model coefficients assigned to cell type one-hot encoding features and scaled remaining ion channel model coefficients.

### Statistical analyses

In general, violin plots depict a symmetric kernel density estimate of the data with an embedded box-and-whisker plot with the white circle denoting median; the box denoting the interquartile range (IQR) between the first and third quantile (Q1 and Q3); and the whiskers extending to at maximum 1.5*IQR above and below Q3 and Q1.

To robustly assess model performance, we generated a distribution of *R*-values by fitting models to 100 different train-test splits of the data. To compare these distributions across different predicted physiology features, data processing conditions (e.g., cell type averaging or not), input gene sets (e.g., ion channels versus synaptic proteins), cell types, and datasets, we used unpaired, two-sided *t*-tests with Bonferroni correction.

For cross cell type and species analyses, we trained and tested on the full population of cells in each category. To estimate a distribution without explicitly choosing different train-test splits, we computed confidence intervals based on 100 bootstrap samples of the training dataset.

## Data availability

This study was conducted using published Patch-Seq datasets from the Allen Institute.^4,16^ Both PatchSeq datasets can be accessed at https://portal.brain-map.org/cell-types/classes/multimodal-characterization under “Neurons in Mouse Primary Visual Cortex” and “Glutamatergic Neuron Types in Layer 2 and Layer 3 of Human Cortex.”

Mouse interneuron electrophysiology data is available through the DANDI archive with accession number, 000020, and the human pyramidal neuron electrophysiology data is available with accession number 000023. Count files for the mouse dataset are provided at https://data.nemoarchive.org/other/AIBS/AIBS_patchseq/transcriptome/scell/SMARTseq/processed/analysis/20200611/. The human dataset will be deposited by the Allen institute once it has cleared procedures at the NeMO archive for restricted datasets.

### Code availability

Custom code will be available at a Github repository (https://github.com/renniek/AIBS-PatchSeq-Analysis) upon publication of the manuscript.

## Acknowledgments

We thank N. Johansen, N. Gouwens, and B. Kalmbach from the Allen Institute for Brain Science for discussions about methodological approaches and preliminary results. We thank C. Plant and S. Smith for helpful feedback on a draft.

## Author Contributions

R.M.K, S.W.L and S.F.O designed the study; R.M.K. performed data analysis; S.F.O and R.M.K. secured funding; R.M.K. wrote the manuscript with comments and reviewing from all authors.

---

**Supplemental Figure S1.**
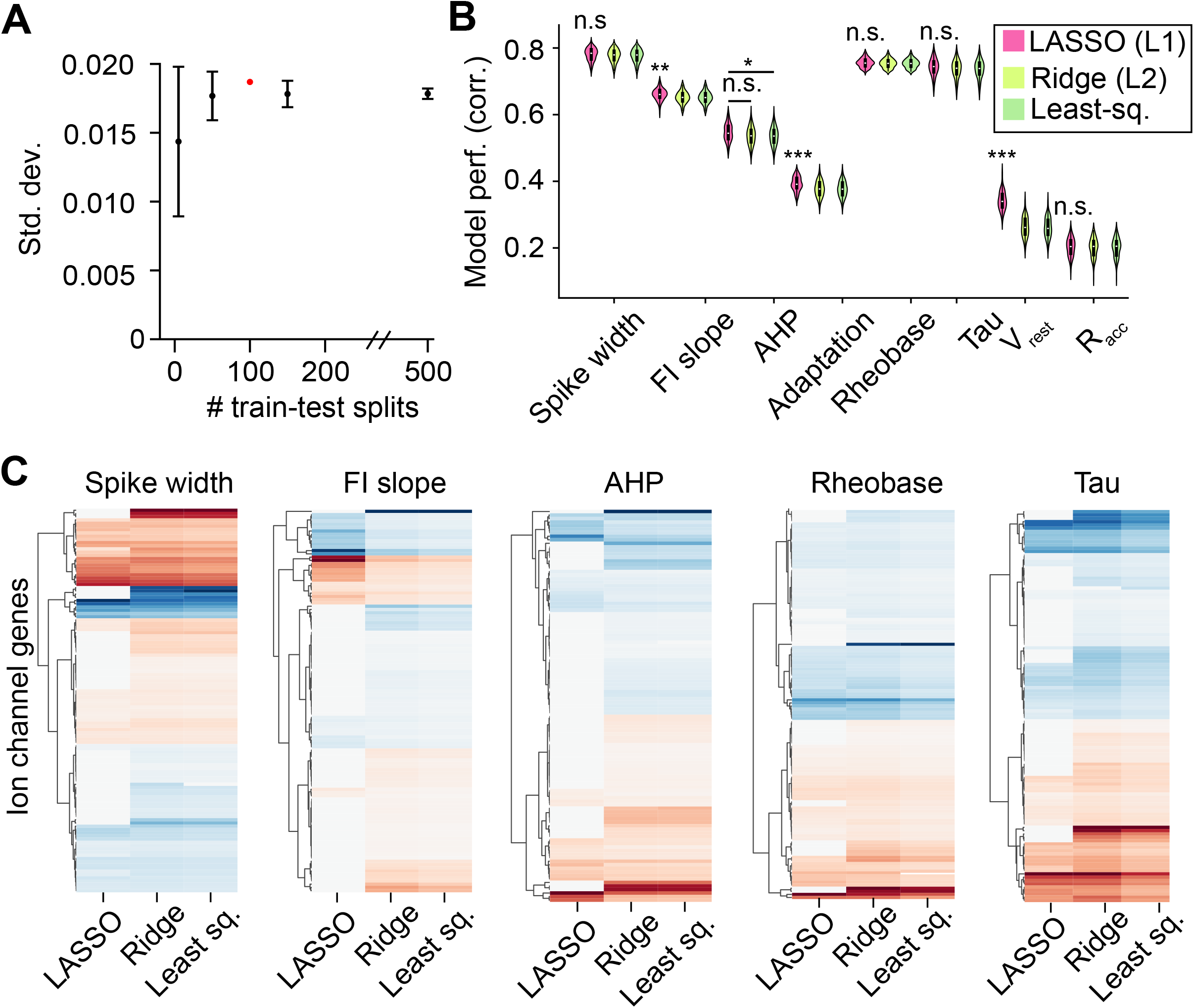
Model validation and comparison. A. Variability in model performance as a function of number of train-test splits. For each number of train-test splits (e.g., 10), we sampled train-test splits 100 times, calculating the standard deviation of model performance across train-test splits for each iteration. The single red-dot captures the variability across the 100 train-test splits used in our analysis, and suggests that additional train-test splits would not capture more variability. **B.** Model performance across different regularization methods. LASSO, or L1-regularization, performed comparably to or statistically significantly better than Ridge (L2-regularization) and least-squares regression. Statistics denote pairwise comparison of all other regression methods to LASSO when specific comparisons are not explicitly noted, and the asterisks denote the following *: *p* < 0.05; **: *p* < 0.005; ***: *p* < 0.001 on unpaired *t*-test with Bonferroni correction. **C.** Model coefficients assigned to ion channel genes for predictive physiology models (mean Pearson’s *R* > 0.5, 100 train-test splits) acros different regular- ization methods. Coefficients were scaled to the absolute maximum value for a given physiology model and genes were clustered according to their assigned coefficients across regularization methods.

**Supplemental Figure S2.**
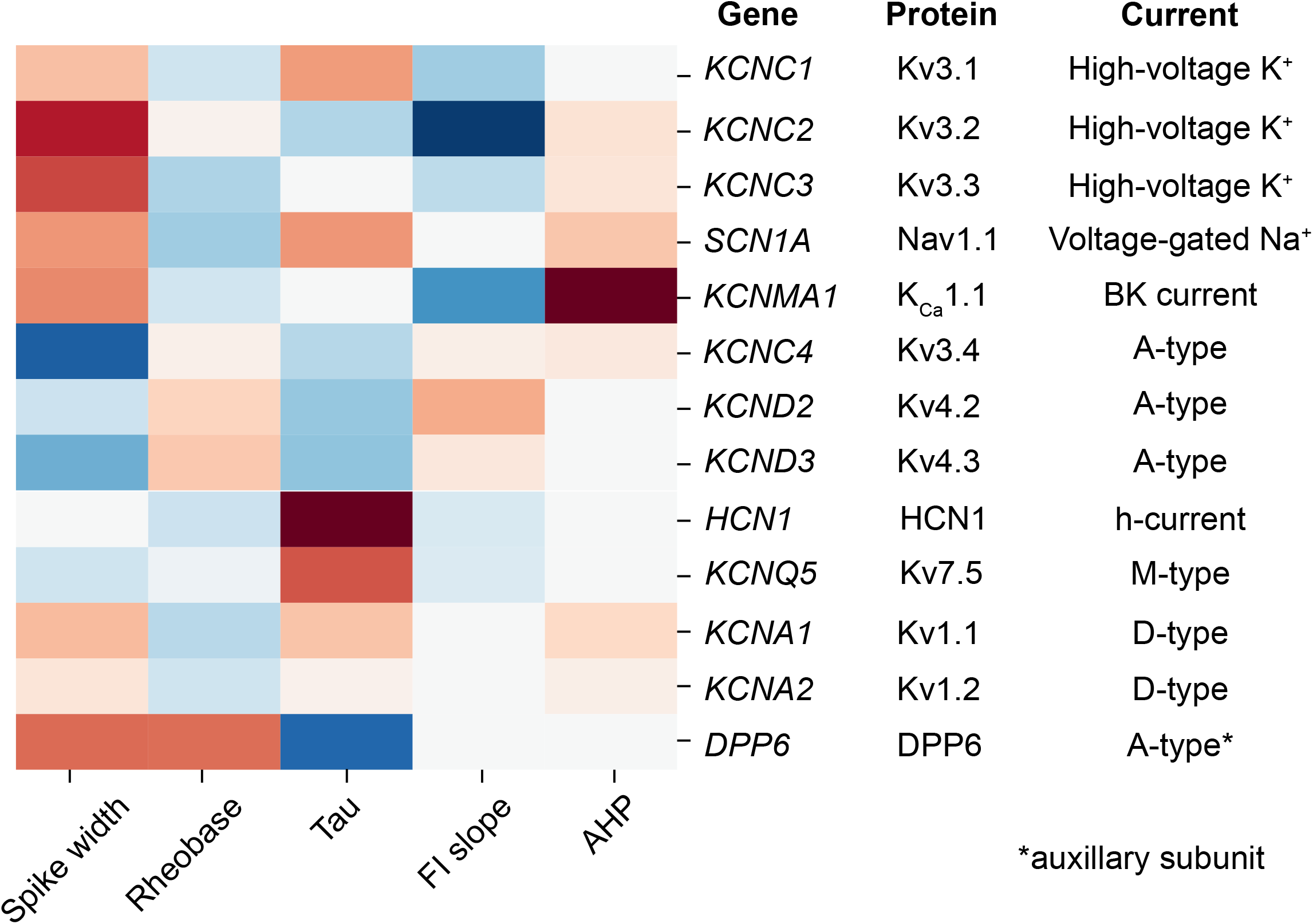
Model coefficients for genes highlighted in the text, and corresponding protein and physiological current. Most genes relate to multiple physiology features, and show heterogeneous relationships to these features gene-to-gene.

**Supplemental Figure S3.**
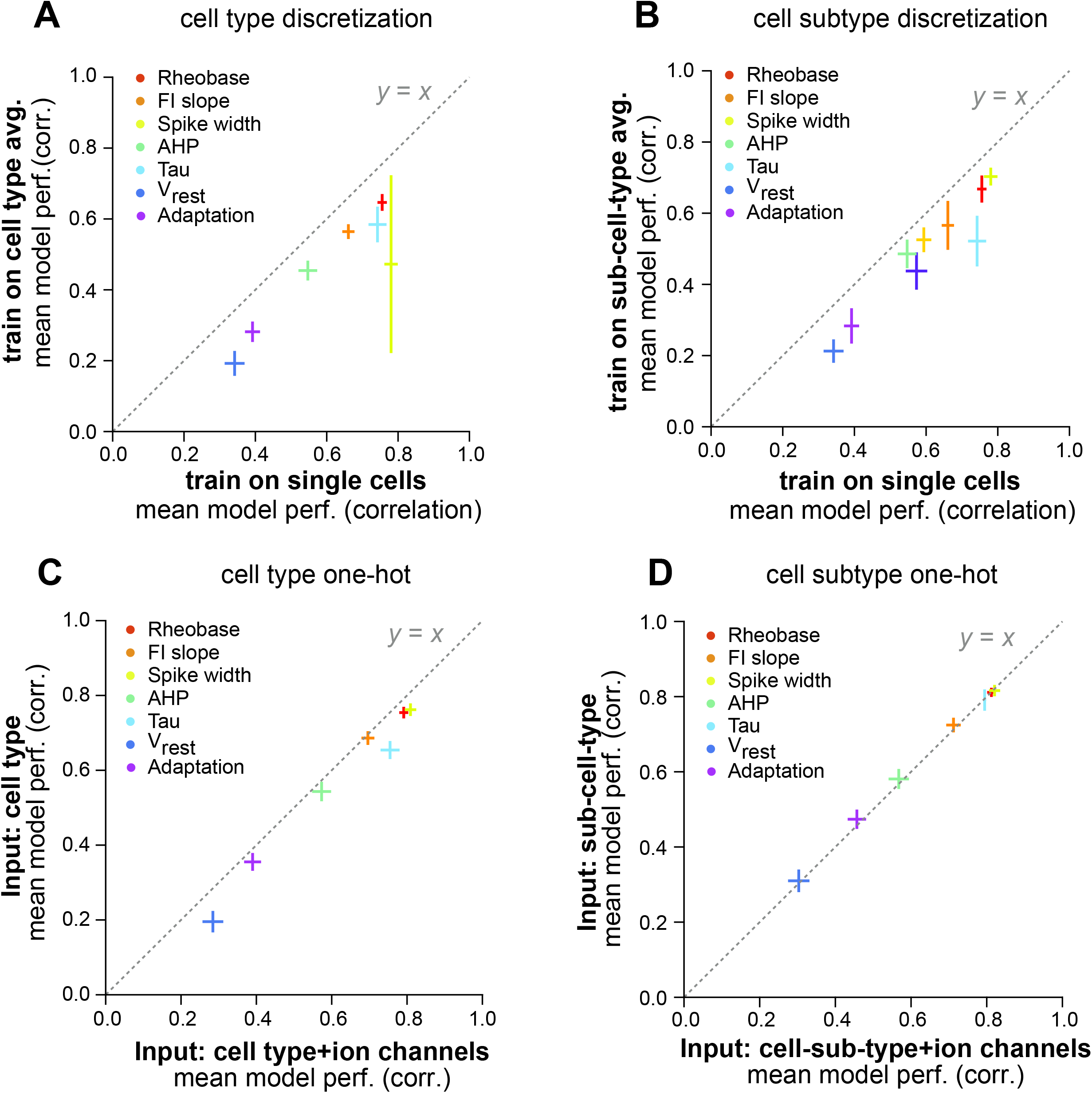
Predictive information in single cells relative to cell types. For all plots, points represent mean correlation between actual versus predicted physiology on held-out samples and error bars the standard deviation of this correlation across 100 different train-test splits of the data. **A.** Training on single-cell gene expression data (*x*-axis) significantly improved model predictions of physiology on held out single cells relative to training on cell type averages (*y*-axis; all *p* < 0.001). **B.** Training on single-cell gene expression (*x*-axis) significantly improved model performance relative to training on cell subtype averages (Sst Tac1 Tacr3; all *p* < 0.001). **C.** Adding ion channel gene expression to information about which cell type each sample belongs to improved model predictive accuracy (*x*-axis versus *y*-axis; all *p* < 0.001). **D.** Adding ion channel gene expression to information about which cell subtype each sample belongs to did not statistically change model performance for Rheobase, Spike Width, tau, and V_rest_ and significantly reduced model performance for FI slope, AHP, and Adaptation with *p* < 0.001 (*x*-axis versus *y*-axis). Pairwise statistical comparisons were made via two-sided, unpaired *t*-tests with Bonferroni correction.

**Supplemental Figure S4.**
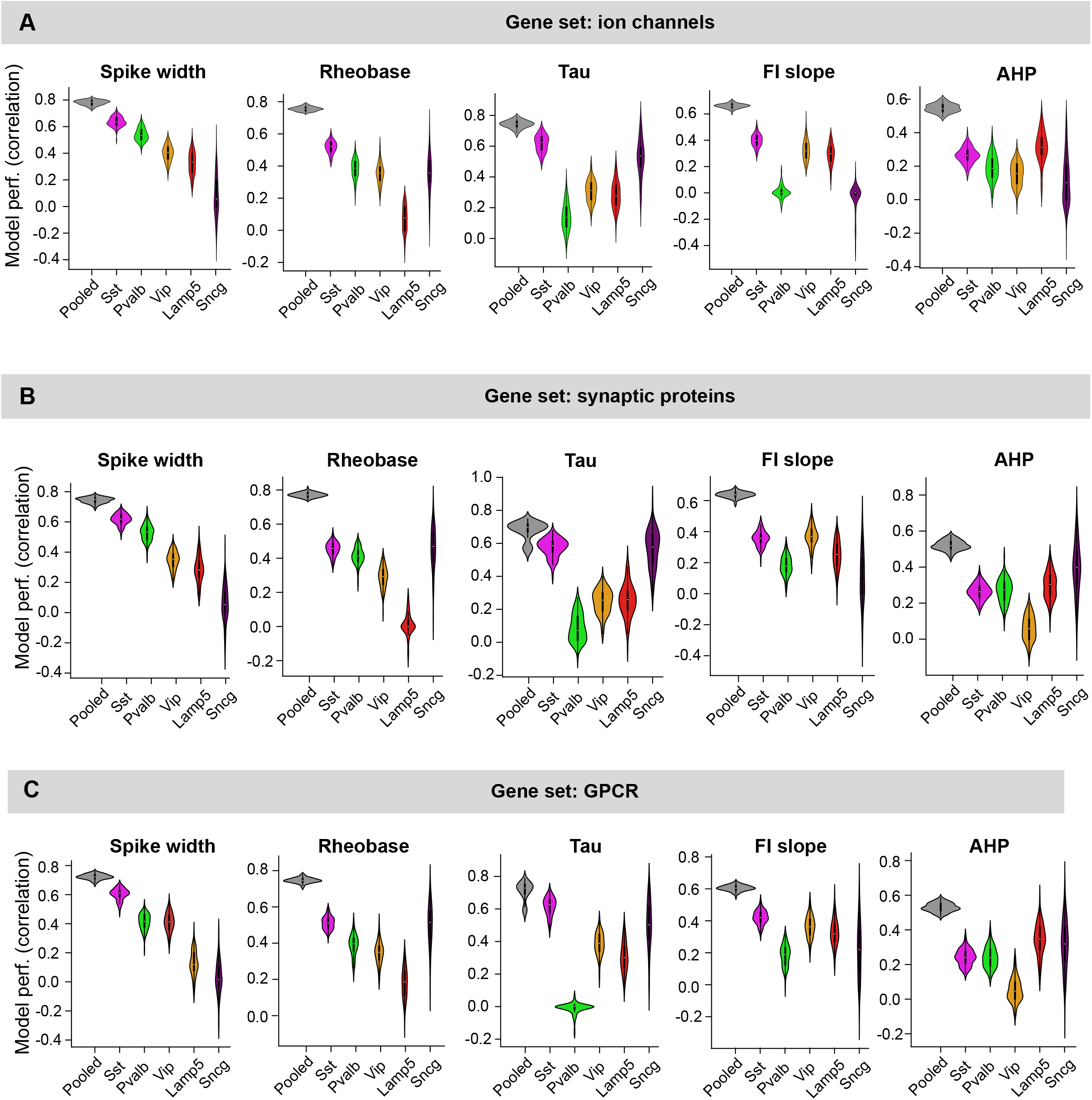
Model performance on pooled dataset versus cell-type-subsetted datasets. A. Violin plots represent correlation between actual versus predicted physiology from ion channel gene expression on held-out test samples for 100 train-test splits. Only physiology features with *R* > 0.5 correlation on pooled dataset predictions shown. **B.** As in (A) but for synaptic protein genes. **C.** As in (A) but for GPCR genes.

**Supplemental Figure S5.**
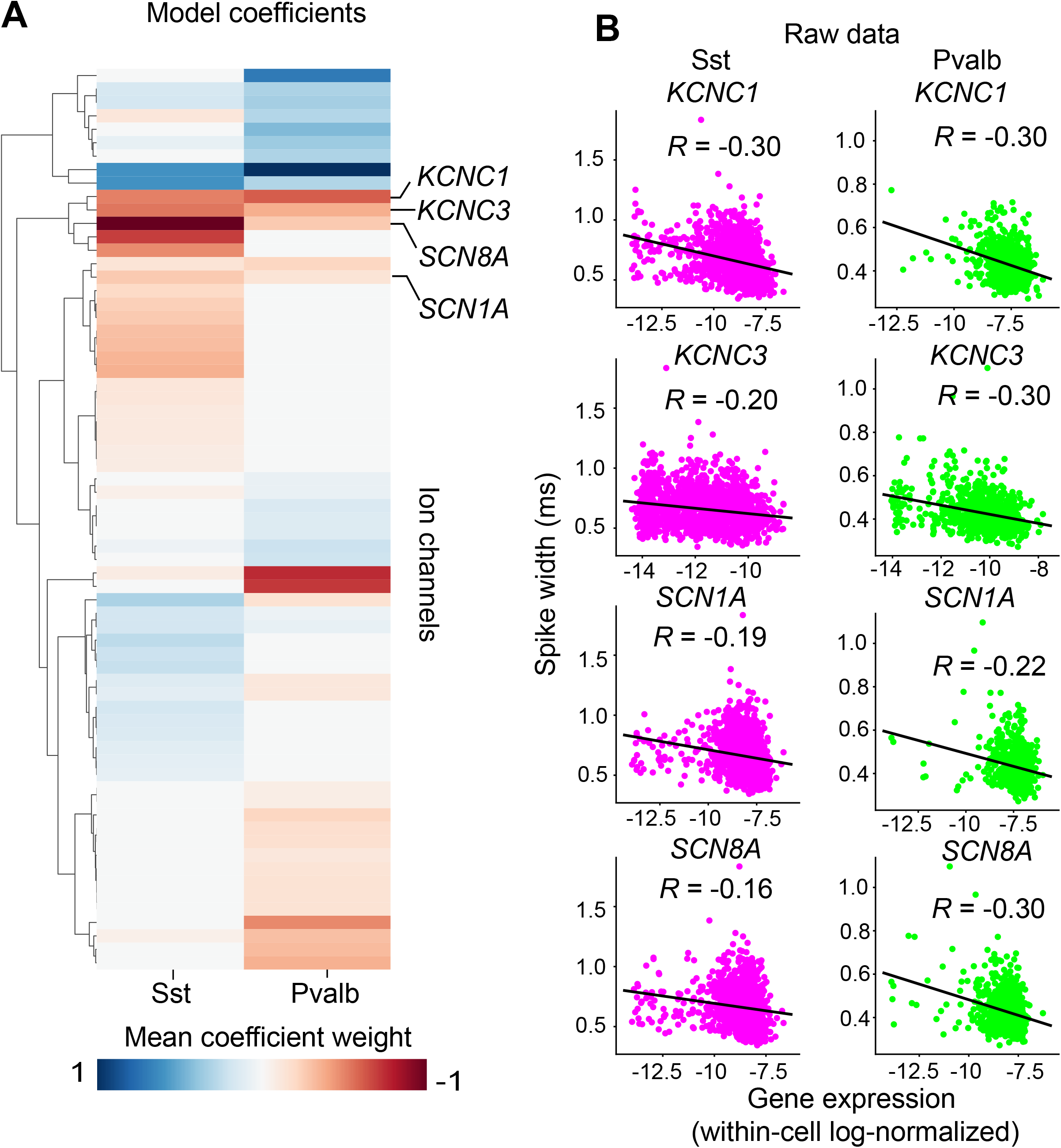
Gene-physiology relationships within Sst and Pvalb interneurons. **A.** Assigned model coefficients for genes with consistent relationships to spike width in within-Sst and within-Pvalb models. **B.** Expression of canonical fast-spiking genes (*KCNC1*, *KCNC3*, *SCN1A*, and *SCN8A*) versus spike width within Sst (*n* = 1463) and Pvalb (*n* = 770) interneurons.

**Supplemental Figure S6.**
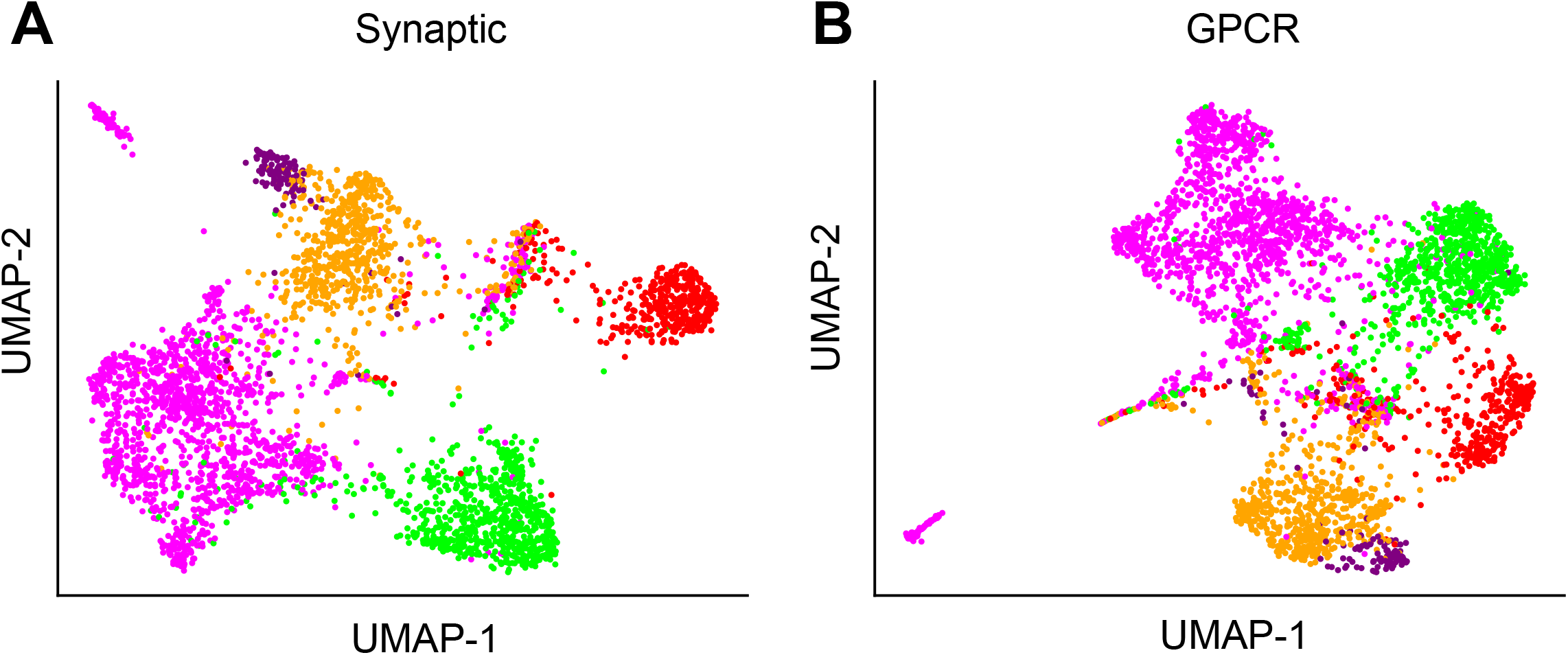
Synaptic and GPCR gene expression delinates canonical interneuron cell types. UMAP on gene expression data as in Figure 2A but for **A.** synaptic and **B.** GPCR genes. For both, *n* = 3651 cells.

**Supplemental Figure S7.**
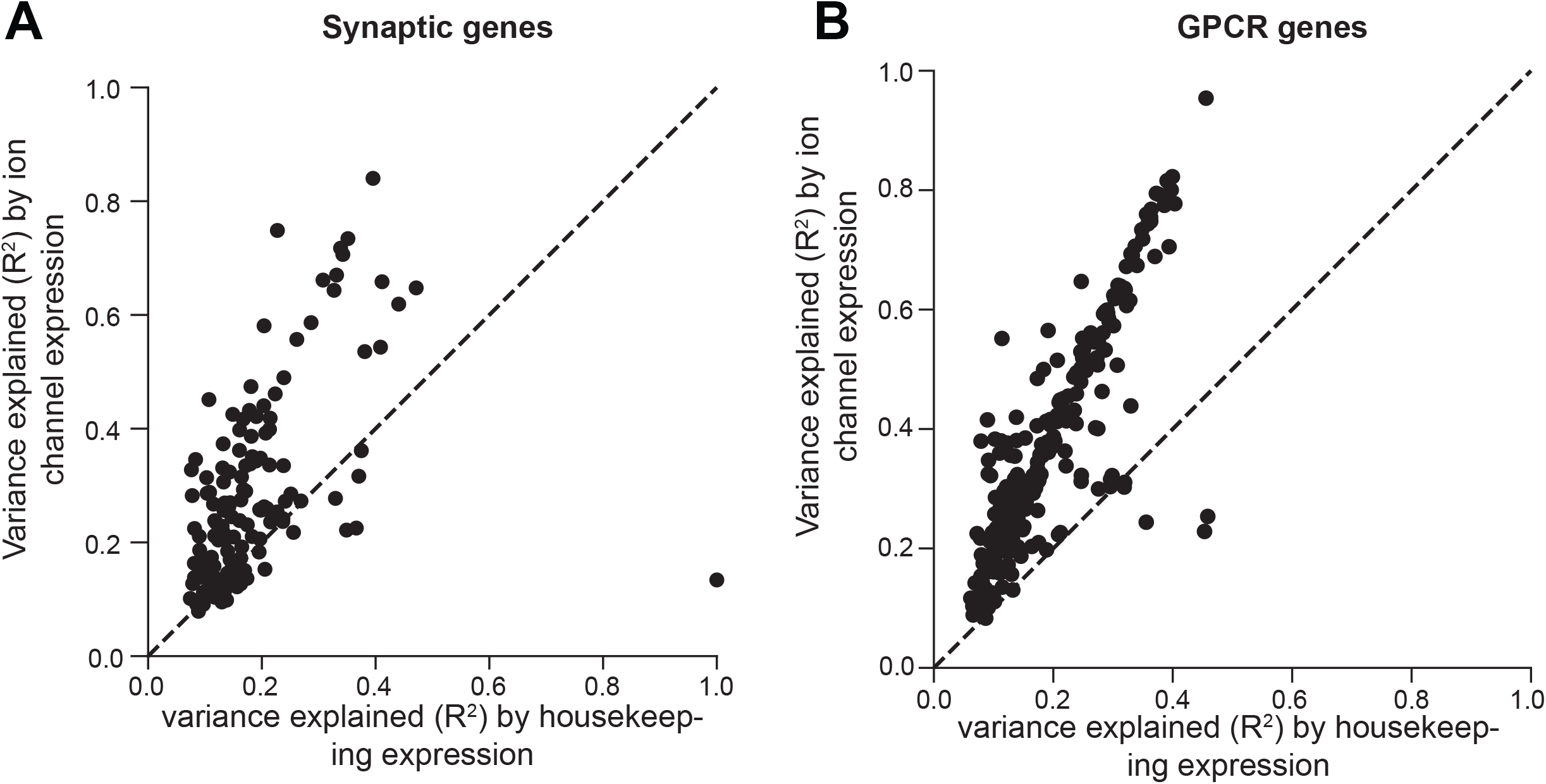
Ion channel gene expression is more predictive of variance in synaptic and GPCR gene expression relative to housekeeping gene expression. A. *R*^2^ (i.e., variance explained) of linear model predictions of synaptic gene expression from housekeeping (*x*-axis) versus ion channel (*y*-axis) gene expression. Each point represents a synaptic gene. **B.** As in A, but for GPCR genes.

**Supplemental Figure S8.**
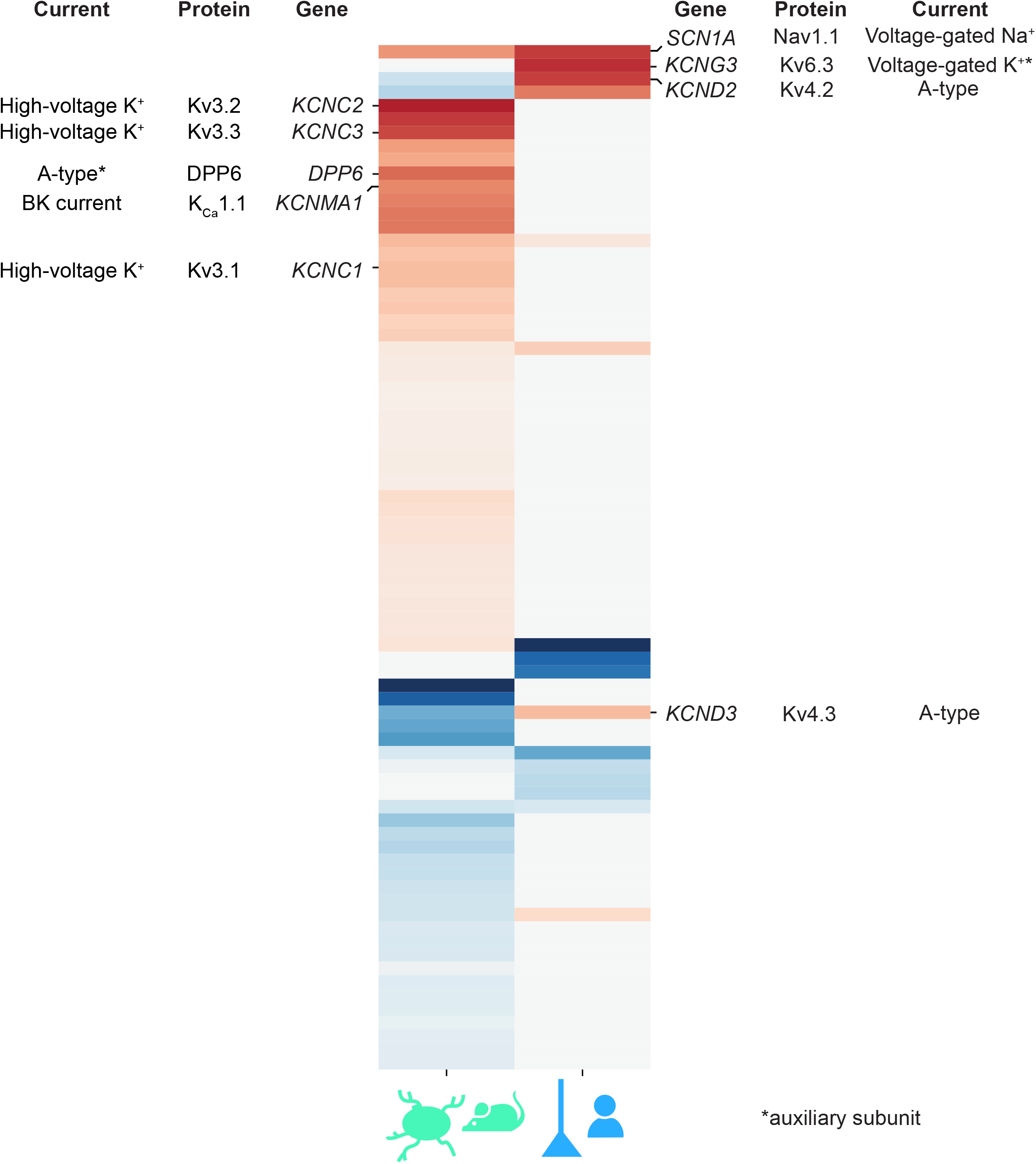
Relationships between spike width and ion channel gene expression are distinctive between mouse interneuron and human pyramidal neurons. Left: Ion channels strongly related to spike width in mouse interneuron dataset. Right: Ion channels strongly related to spike width in human pyramidal neuron dataset. Negative correlations between *SCN1A* and spike width is one of the only conserved relationships identified by our models. Relationships between *KCND2*/*KCND3* and spike width are opposite across datasets.

**Supplemental Table S1. Sample sizes by dataset, cell type, and physiology feature.**

**Supplemental Table S2. Model coefficients for pooled mouse interneuron dataset across physiology feature models and gene sets.** Average cell type coefficients for physiology feature models with mean *R* > 0.5 across train test splits for mouse interneuron dataset. Genes with coefficients with consistent non-zero sign across 80% of train-test splits displayed, and coefficient values are averaged across 100 train-test splits.

**Supplemental Table S3. Model coefficients missed by LASSO.** Genes whose model coefficients were dropped to 0 by LASSO for a given physiology model and had non-zero coefficients assigned by ridge and least-squares. As before, coefficients were filtered such that only coefficients with consistent relationships across 0.8 of 100 train-test splits are displayed, and their averaged value across those 100 train-test splits are displayed.

**Supplemental Table S4. Model coefficients for relating ion channel gene expression to spike width across different training set formats.** Training sets comprised of: A) cell type-averaged ion channel gene expression and spike width data (Cell type avg ion channel); B) single-cell ion channel gene expression and categorical variable that denotes cell type (Cell type+ion channels); C) and D) as in (A) and (B) but for cell-sub-type; E) single-cell ion channel gene expression and spike width data. As before, average cell type coefficients for physiology feature models with mean *R* > 0.5 across train test splits for mouse interneuron dataset. Genes with coefficients with consistent non-zero sign across 80% of train-test splits displayed, and coefficient values are averaged across 100 train-test splits. As before, coefficients were filtered such that only coefficients with consistent relationships across 0.8 of 100 train-test splits are displayed, and their averaged value across those 100 train-test splits are displayed.

**Supplemental Table S5. Average model coefficients for within-Sst and Pvalb models for mouse interneuron dataset.** Genes with coefficients with consistent non-zero sign across 80% of train-test splits displayed, and coefficient values are averaged across 100 train-test splits. Models highlighted in text shown for different gene sets and physiology features.

**Supplemental Table S6. Model coefficients for human pyramidal neuron dataset for spike width model.** Average cell type coefficients for spike width model, which had mean *R* > 0.5 across train test splits for human pyramidal neuron dataset. Genes with coefficients with consistent non-zero sign across 80% of train-test splits displayed, and coefficient values are averaged across 100 train-test splits.

Supplemental Table S8. Metadata and pooled physiology feature dataset for mouse interneuron and human pyramidal neuron dataset.

**Supplemental Table S9. Model coefficients for mouse interneuron vs. human pyramidal neuron models predicting spike width from ion channel gene expression.** Average cell type coefficients for spike width model, which had mean *R* > 0.5 across train test splits for human pyramidal neuron dataset. Genes with coefficients with consistent non-zero sign across 80% of train-test splits displayed, and coefficient values are averaged across 100 train-test splits.

